# Dietary Monoterpenoids As a New Class of Allosteric Human Aryl Hydrocarbon Receptor Antagonists

**DOI:** 10.1101/2020.11.30.404178

**Authors:** Karolína Poulíková, Iveta Zůvalová, Barbora Vyhlídalová, Kristýna Krasulová, Eva Jiskrová, Radim Vrzal, Sandhya Kortagere, Martina Kopečná, David Kopečný, Marek Šebela, Katharina Maria Rolfes, Thomas Haarmann-Stemmann, Sridhar Mani, Zdeněk Dvořák

**Affiliations:** Department of Cell Biology and Genetics, Faculty of Science, Palacký University, Šlechtitelů 27, 783 71 Olomouc, Czech Republic; Department of Protein Biochemistry and Proteomics, Centre of the Region Hana, Faculty of Science, Palacký University, Šlechtitelů 27, 783 71 Olomouc, Czech Republic; Department of Genetics and Department of Medicine, Albert Einstein College of Medicine, Bronx, NY 10461, U.S.A; Department of Microbiology & Immunology, Drexel University College of Medicine, Philadelphia, PA 19129, U.S.A; IUF-Leibniz-Research Institute for Environmental Medicine, 40225, Düsseldorf, Germany

**Author notes:** **Corresponding authors:** Zdeněk Dvořák, Department of Cell Biology and Genetics Faculty of Science, Palacky University Olomouc Slechtitelu 27; 783 71 Olomouc; Czech Republic, E, Sridhar Mani Department of Genetics and Department of Medicine Albert Einstein College of Medicine Bronx, NY 10461, U.S.A., E.

## Abstract

Carvones, the constituents of essential oils of dill, caraway, and spearmint, were reported to antagonize the human aryl hydrocarbon receptor (AhR); however, the exact molecular mechanism remains elusive. We show that carvones are non-competitive allosteric antagonists of the AhR that inhibit the induction of AhR target genes in a ligand-selective and cell type-specific manner. Carvones do not displace radiolabeled ligand from binding at the AhR, but they bind allosterically within the bHLH/PAS-A region of the AhR. Carvones did not influence a translocation of ligand-activated AhR into the nucleus. Carvones inhibited the heterodimerization of the AhR with its canonical partner ARNT and subsequent binding of the AhR to the promotor of CYP1A1. Interaction of carvones with potential off-targets, including ARNT and protein kinases, was refuted. This is the first report of a small dietary monoterpenoids as a new class of AhR non-competitive allosteric antagonists with the potential preventive and therapeutic application.

## INTRODUCTION

The aryl hydrocarbon receptor (AhR) is a ligand-activated transcription factor that belongs to the family of basic helix-loop-helix transcription factors. In the resting state, unliganded AhR resides in the cytosol. Upon the ligand binding to the AhR, the complex ligand-receptor translocates to the cell nucleus. It forms a heterodimer with AhR nuclear translocator (ARNT), which binds to the specific response elements in the target genes’ promoters. Typical xenobiotic ligands of the AhR are environmental contaminants such as polyaromatic hydrocarbons (e.g., benzo[a]pyrene - BaP) and halogenated aromatic hydrocarbons (e.g., 2,3,7,8-tetrachlorodibenzo-*p*-dioxin - TCDD), but also naturally occurring chemicals such as various polyphenols. Endogenous ligands of AhR are mainly intermediary and microbial metabolites of tryptophan, such as 6-formylindolo[3,2-b]carbazole (FICZ) [1]. The AhR regulates the expression of genes involved in xenoprotection, immune response, cell cycle, differentiation, lipid, and carbohydrate metabolism. Thereby, AhR is a pivotal determinant not only in human physiology (e.g., hematopoietic development)[2] but also in the incidence, onset, and progress of many pathophysiological processes, including carcinogenesis, inflammation, infection, diabetes, and cardiovascular diseases [3,4].

New selective AhR ligands’ development has received attention in recent years because of their potential therapeutic and preventive potential [5,6]. For instance, rational drug design that included screening a chemical library of indoles and indazoles resulted in developing small molecules PY109 and PY108, which are highly potent AhR agonists (EC_50_ ~1.2 nM). These drug-like indole mimics has demonstrated anti-inflammatory properties in mice, where they potently induced IL-22 and expanded tissue ILC3 and γδ T cell subpopulations [7]. Also, repositioning of clinically used AhR-active drugs such as tranilast, flutamide, or omeprazole was proposed as AhR-dependent chemotherapy to treat breast and pancreatic cancers [8]. The drawback with all these compounds for long term use are side effects and off-target effects of the drugs [9].

It is worthy of pointing out that most AhR ligands are partial agonists, which dose-dependently activate the AhR and at the same time behave as competitive antagonists when applied simultaneously with another, usually potent AhR agonist. Pure agonists of the AhR are, for example, highly potent and efficacious ligands such as TCDD, whereas pure antagonists are scarce. For instance, stilbenoid resveratrol or synthetic inhibitor of c-Jun-N-terminal kinase SP600125 had a long time been deemed as the AhR antagonists until their minimal residual agonist activity was unveiled [10]. The first identified and *bona fide* frequently used, the pure antagonist of the AhR was 3’-methoxy-4’-nitroflavone (MNF), which bound with high affinity (K_i_ ~1.5 nM) at rat hepatic cytosolic AhR, and competitively displaced ^3^H-TCDD [11]. However, several studies reported that AhR-dependent enhanced CYP1A1 transcription by MNF [12]. By a screening of a chemical library composed of 10,000 compounds, 2-methyl-2*H*-pyrazole-3-carboxylic acid (2-methyl-4-*o*-tolylazo-phenyl)-amide (CH223191) was identified as a potent antagonist (IC_50_ ~ 30 nM) of the AhR. It displaced TCDD from binding at the mouse AhR; thereby, the action mechanism was competitive [13]. Whereas a series of CH223191-based antagonists were developed, later on, the AhR-independent pro-proliferative properties of CH223191 were reported [14]. Also, CH223191 is a ligand-selective antagonist of the AhR. CH223191 preferentially inhibits halogenated aromatic hydrocarbons class of agonists (e.g., TCDD), but not others, like polyaromatic hydrocarbons or flavonoids [15]. The Perdew lab reported *N*-(2-(1*H*-indol-3-yl)ethyl)-9-isopropyl-2-(5-methyl pyridine-3-yl)-9*H*-purin-6-amine (GNF351) as high affinity (IC_50_ ~ 62 nM) pure competitive antagonist of the AhR with a capability to inhibit both genomic and non-genomic actions of the AhR [16]. Given low intestinal absorption and extensive metabolism following an oral administration, GNF351 was proposed as a site-specific antagonist of the AhR in the intestine and colon [5]. There are isolated reports on the *in vitro* and *in vivo* effects of FDA-approved drugs with AhR-antagonist activity. For instance, clofazimine, an anti-leprosy drug and AhR antagonist, suppressed multiple myeloma in transgenic mice; however, the putative AhR-dependent mechanism was not directly evidenced [17]. Another example is the relapse during melanoma treatment with BRAF inhibitor vemurafenib, which was suggested to be delayed by targeting the constitutively active AhR in persisting cells with antagonists [18]. Moreover, we have identified vemurafenib as the competitive antagonist of the AhR, which inhibited *in vitro* and *in vivo* effects of AhR-dependent processes, including the abrogation of anti-inflammatory signaling and response [19].

We recently reported that the essential oils of dill, caraway, and spearmint have antagonist effects on the AhR and that carvones, which are the major constituents of these oils, are responsible for the AhR antagonism [20]. Carvone is a monocyclic monoterpenoid in two optical conformers, “S” and “R”. Humans distinguish the odor quality between sweetish, spearmint-like R-carvone and spicy, caraway-like S-carvone. Both carvones activate odorant receptor OR1A1 but displaying selectivity for individual enantiomers [21]. While we showed that carvones are antagonists of the AhR, the exact mechanism of how they influence the AhR signaling pathway remains elusive. Humans’ exposure to carvones occurs mainly through the dietary intake of carvones-containing foods and beverages [20] and *via* percutaneous absorption because carvones are used as skin permeabilizers in transdermal patches [22]. In the current study, we investigated in detail the antagonist effects of carvones on the human AhR, and we aimed to decipher their molecular mechanism of action. We describe the atypical, allosteric, and non-competitive mechanism of AhR antagonism, involving disruption of AhR-ARNT dimerization by carvones in the cell nucleus. This is the first report that small dietary monoterpenoids are a new class of AhR non-competitive allosteric antagonists with the potential preventive and therapeutic application.

## METHODS

### Chemicals and materials

S-carvone (sc-239480, purity 99.4 %, Lot L0613), R-carvone (sc-293985, purity 99.7 %, Lot H1015), and D-limonene (sc-205283, Lot F1314) were purchased from Santa Cruz Biotechnology. BaP (B1760, Lot SLBS0038V, purity 99 %), FICZ (SML1489, Lot 0000026018, purity 99.5 %), staurosporine (S4400, purity 98%), deferoxamine mesylate (DFX; D9533, purity 92.5%) and dexamethasone (DEX; D4902, Lot 112K12845, purity 98 %) were obtained from Sigma-Aldrich. TCDD (RPE-029) was purchased from Ultra Scientific, and 2,3,7,8-tetrachlorodibenzofuran (TCDF; Amb17620425, Lot 51207-31-9) was obtained from Ambinter. Radio-labelled [^3^H]-TCDD (ART 1642, Lot 181018, purity 98.6 %) was purchased from American Radiolabeled Chemicals. Bio-Gel® HTP Hydroxyapatite (1300420, Lot 64079675) was obtained from Bio-Rad Laboratories.

### Cell lines and hepatocytes

Human hepatoma cells HepG2 (ECACC No. 85011430), intestinal human colon adenocarcinoma cells LS180 (ECACC No. 87021202), human immortalized keratinocytes HaCaT (kindly donated by P. Boukamp, IUF Düsseldorf, Germany), and mouse hepatoma Hepa1c1 (ECACC No. 95090613) were cultured as recommended by the supplier. Primary human hepatocytes LH75 (female, 78 years, Caucasian) were prepared at the Faculty of Medicine, Palacky University Olomouc. The tissue acquisition protocol complied with the regulation issued by “Ethical Committee of the Faculty Hospital Olomouc, Czech Republic” and Transplantation law #285/2002 Coll. Primary human hepatocytes Hep200571 (male, 77 years, unknown ethnicity) and Hep220993 (female, 76 years, Caucasian) were purchased from Biopredic International (Rennes, France). Mycoplasma Detection Kit-Digital Test v2.0 Cat.No. B39132 (Biotool) was used to survey mycoplasma infection.

### Reporter gene assays

The stably transfected gene reporter cell line AZ-AHR, derived from human hepatoma cells HepG2, expressing endogenous AhR and transfected with a construct containing several AhR binding sites upstream of a luciferase reporter gene, was used for the evaluation of the transcriptional activity of the AhR [23]. Cells were seeded at 96-well culture plates, and following 16 h of stabilization, they were incubated for 24 h with tested compounds or their combinations. After that, the cells were lysed, and luciferase activity was measured on a Tecan Infinite M200 Pro plate reader (Schoeller Instruments, Czech Republic). Half-maximal inhibitory concentrations (IC_50_), half-maximal effective concentrations (EC_50_), and concentrations of EC_80_ were calculated using GraphPad Prism 8 software (GraphPad Software, San Diego, U.S.A.). Experiments were performed in at least two independent cell passages. Incubations and measurements were performed in quadruplicates (i.e., four technical replicates). (ref. Data in Figure 1).

**Figure 1.**
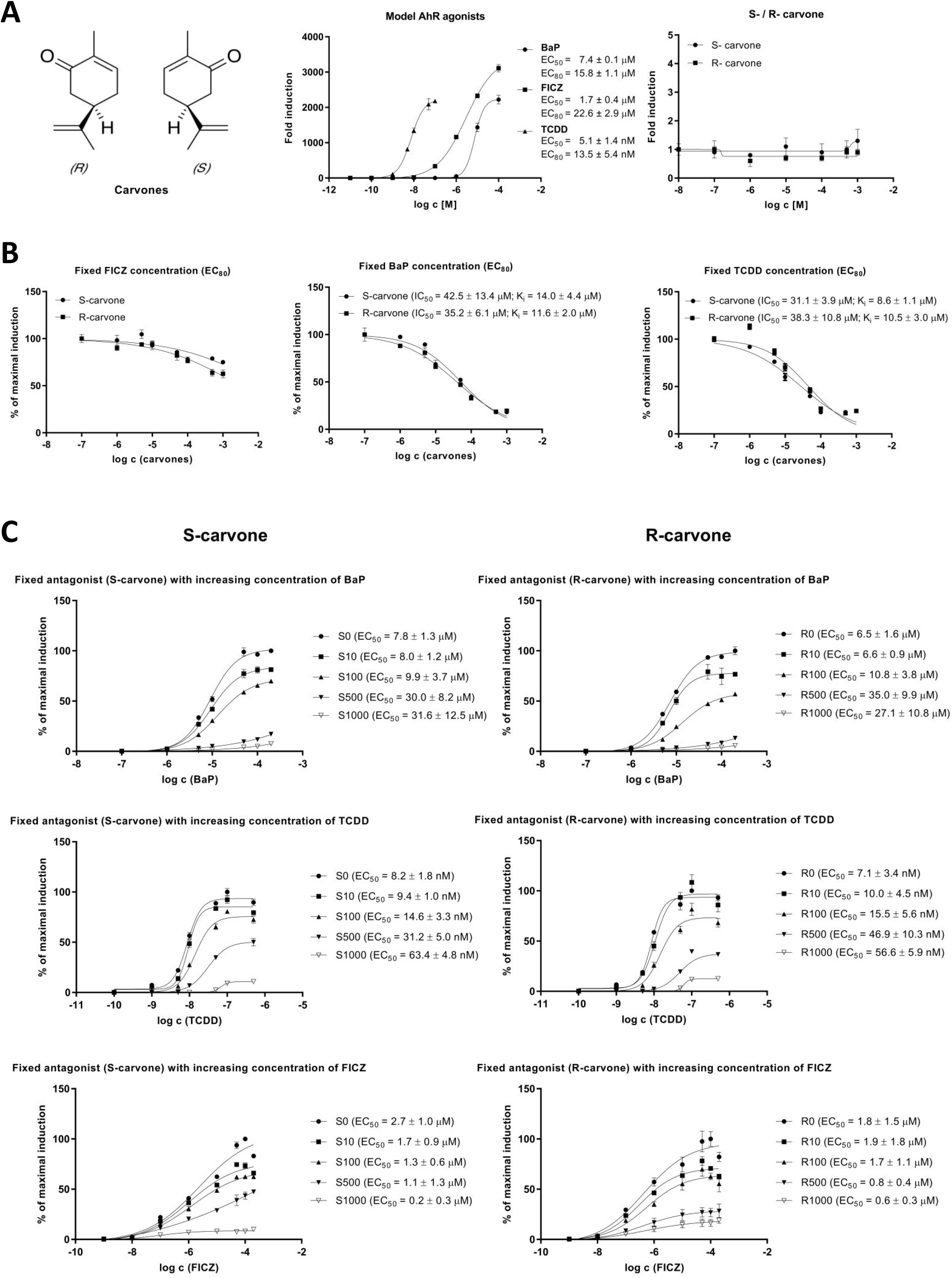
Non-competitive antagonism of the AhR by carvones. Reporter gene assay was carried out in stably transfected AZ-AHR cells, incubated for 24 h with tested compounds. Experiments were performed in two independent cell passages. Incubations and measurements were performed in quadruplicates (technical replicates). **(A)** Basal activity of AhR: left – the structure of carvones; middle - effects of AhR agonists TCDD, BaP, and FICZ (EC_50_ and EC_80_ values indicated in the graph); right – effects of carvones. **(B)** Agonist-inducible AhR activity: combined incubations with fixed concentrations of agonists (at EC_80_) and increasing concentrations of carvones (IC_50_ and K_i_ values indicated in the graph). **(C)** Non-competitive antagonism of carvones: combined incubations with a fixed concentration of carvones and increasing concentrations of AhR agonists.

### Quantitative real-time polymerase chain reaction qRT-PCR

The total RNA was isolated using TRI Reagent® (Sigma-Aldrich). cDNA was synthesized from 1 μg of total RNA using M-MuLV Reverse Transcriptase and Random Primers 6 (both New England Biolabs) at 42 °C for 60 min and diluted in 1:4 ratio by PCR grade water. qRT-PCR was carried out on Light Cycler® 480 Instrument II (Roche). Data were processed by the delta-delta C_t_ method and normalized *per GAPDH* as a housekeeping gene. The levels of *GAPDH* and *CYP1A1* mRNAs were determined using probes and primers from Universal Probes Library (UPL; Roche). GAPDH–UPL60, fw: CTCTGCTCCTCCTGTTCGAC, rev: ACGACCAAATCCGTTGACTC and CYP1A1– UPL33, fw: CCAGGCTCCAAGAGTCCA, rev: GATCTTGGAGGTGGCTGCT. Eurofins Genomics primers were used for *VEGF* mRNA (vascular endothelial growth factor), fw: TGCAAAAACACAGACTCGCG, rev: TGTCACATCTGCAAGTACGTTCG; and *GAPDH* mRNA, fw: AGGTGAAGGTCGGAGTCA, rev: GGTCATTGATGGCAACAA.

### Simple western blotting by Sally Sue™

Total protein extract was isolated by using ice-cold lysis buffer (150 mM NaCl; 10 mM Tris pH 7.2; 1% (v/v) Triton X-100; 0.1% (w/v) SDS; 1% (v/v) sodium deoxycholate; 5 mM EDTA; anti-protease cocktail; anti-phosphatase cocktail) and protein concentration was determined using Bradford reagent. Detection of CYP1A1 and β-actin proteins was performed by Sally Sue™ Simple Western System (ProteinSimple™) using the Compass Software version 2.6.5.0 (ProteinSimple™). Immuno-detection was performed using a primary antibody against CYP1A1 (mouse monoclonal, sc-393979, A-9, dilution 1:100, Santa Cruz Biotechnology) and β-actin (mouse monoclonal, 3700S, dilution 1:100, Cell Signalling Technology). Detection was performed by horseradish-conjugated secondary antibody followed by reaction with a chemiluminescent substrate.

### 7-ethoxyresorufin-*O*-deethylase activity (EROD)

AZ-AhR cells plated at 96-well culture dishes were incubated for 24 hours with vehicle (DMSO; 0.1% v/v), TCDD (13.5 nM) and/or S-carvone (1 mM)+TCDD (13.5 nM). After washing with PBS, the medium containing 7-ethoxyresorufin (8 μM) and dicoumarol (10 μM) was applied to the cells. Culture plates were incubated at 37 °C for 30 min. After that, an aliquot of 75 μl of the medium was mixed with 125 μl of methanol, and fluorescence was measured in a 96-well plate with 530 nm excitation and 590 nm emission filters, using Tecan Infinite M200 Pro plate reader (Schoeller Instruments, Czech Republic).

### Radioligand binding assay

Cytosol from murine hepatoma Hepa1c1c7 cells was isolated as described [24]. Cytosolic protein (2 mg/mL) was incubated for 2 h at room temperature in the presence of 2 nM [^3^H]-TCDD with S-carvone (1 μM, 10 μM, 100 μM, 1000 μM), FICZ (10 nM; positive control), DEX (100 nM; negative control) or vehicle (DMSO; 0.1% V/V; corresponds to *specific binding of [^3^H]-TCDD = 100%*). Ligand binding to the cytosolic proteins was determined by the hydroxyapatite binding protocol and scintillation counting. Specific binding of [^3^H]-TCDD was determined as a difference between total and non-specific (TCDF; 200 nM) reactions. Five independent experiments were performed, and the incubations and measurements were done in triplicates in each experiment (technical replicates).

### Intracellular distribution of AhR

Immunofluorescence assay was performed as recently described [25]. Briefly, LS180 cells were seeded on chamber slides (ibidi GmbH, Germany) and cultured for two days. Then, cells were treated for 90 min with carvones (1000 μM) in combination with vehicle (0.1% DMSO) or the AhR agonists TCDD (20 nM), BaP (7 μM), and FICZ (8 nM). After the treatment, washing, fixation, permeabilization, and blocking, cells were incubated with Alexa Fluor 488 labeled primary antibody against the AhR (sc-133088, Santa Cruz Biotechnology, U.S.A.) diluted 1:500 in 0.5% bovine serum albumin at 4 °C overnight. The next day, nuclei were stained by 4’,6-diamino-2-phenylindole (DAPI), and cells were enclosed by VectaShield® Antifade Mounting Medium (Vector Laboratories Inc., U.S.A.). The AhR translocation into the nucleus was visualized and evaluated using fluorescence microscope IX73 (Olympus, Japan). The whole staining protocol was performed in two independent experiments in technical duplicates (with all tested compounds). The AhR translocation was evaluated visually depending on the distinct signal intensity of the AhR antibody in the nucleus and cytosol. For percentage calculation, approximately one hundred cells from at least four randomly selected fields of view in each replicate were used.

### Protein immunoprecipitation assay

Effects of carvones on ligand-dependent hetero-dimerization of the AhR with ARNT were studied in cell lysates from LS180 cells incubated with carvones (1000 μM) in combination with vehicle (0.1% DMSO) or the AhR agonists TCDD (20 nM), BaP (7 μM) and FICZ (8 nM) for 90 min at 37°C. Pierce™ Co-Immunoprecipitation Kit (Thermo Fisher Scientific) was used. In brief, 25 μg of AhR antibody (mouse monoclonal, sc-133088, A-3, Santa Cruz Biotechnology) was covalently coupled on resin for 120 minutes at room temperature. The antibody-coupled resin was incubated with cell lysate overnight at 4 °C. In parallel with total parental lysates, eluted protein complexes were diluted in delivered sample buffer and resolved on 8% SDS-PAGE gels followed by Western blot analysis and immuno-detection with ARNT 1 antibody (mouse monoclonal, sc-17812, G-3, Santa Cruz Biotechnology). Chemiluminescent detection was performed using horseradish peroxidase-conjugated anti-mouse secondary antibody (7076S, Cell Signaling Technology) and WesternSure® PREMIUM Chemiluminescent Substrate (LI-COR Biotechnology) by C-DiGit® Blot Scanner (LI-COR Biotechnology).

### Chromatin immunoprecipitation assay

The assay was performed as recently described [25]. Briefly, HepG2 cells were seeded in a 60-mm dish, and the following day they were incubated with carvones (1000 μM) in combination with vehicle (0.1% DMSO) or the AhR agonists TCDD (20 nM), BaP (7 μM), and FICZ (8 nM) for 90 min at 37°C. The procedure followed the manufacturer’s recommendations for SimpleChIP Plus Enzymatic Chromatin IP kit (Magnetic Beads) (Cell Signaling Technology; #9005). Anti-AhR rabbit monoclonal antibody was from Cell Signaling Technology (D5S6H; #83200). CYP1A1 promoter primers were: (fw:AGCTAGGCCATGCCAAAT, rev: AAGGGTCTAGGTCTGCGTGT-3’). Experiments were performed in three consecutive cell passages.

### Protein kinase C inhibition assay

Protein kinase C (PKC) inhibition was assayed in lysates from HepaG2 cells, using PKC Kinase Activity Assay Kit (ab139437; Abcam). Cells were grown to 90% confluency in a 60 mm dish. After removing the medium, 1 mL of lysis buffer (E4030, Promega) was applied for 10 minutes on ice. Cells were scraped, sonicated, and centrifuged at 13 000 rpm/15 minutes/4°C (Eppendorf Centrifuge 5415R; Eppendorf, Stevenage, U.K.). Then, 3 μL of cell lysate were mixed with 297 μL of Kinase Assay Buffer and aliquoted 40 μL into 0.5 mL microtubes. These aliquots were mixed with 1/100 stock solutions of carvones, resulting in final concentrations of 10 μM, 100 μM, and 1000 μM. DMSO (1% V/V) and staurosporine (1 μM) were negative and positive control, respectively. The reaction was initiated by the addition of 10 μL of reconstituted ATP, and the rest of the procedure was performed as described in the manufacturer’s recommendation. Absorbance was measured at 450 nm using microplate reader Infinite M200 (TECAN, Austria). Results are expressed as % of the negative control. The cell lysate was stored at −80°C and used for performing three independent experiments.

### KINOMEscan™ profiling

The KINOMEscan™ screening platform (scanMAX assay) employs a proprietary active site-directed competition binding assay that quantitatively measures interaction between test compounds (here 100 μM S-carvone) and 468 human protein kinases [26]. The assay was performed at Eurofins DiscoverX (San Diego, CA, U.S.A.).

### Molecular modeling and Docking

The crystal structure complex of a construct of the human AhR with a truncated mouse ARNT has been solved (PDB code: 5NJ8) [27]. Since the solved structure does not contain the LBD of the AhR, it was modeled based on neuronal PAS-1 protein (pdb code: 5SY5) [28] as previously described [25]. The molecular structures of carvones were modeled using the ligand builder module of the Molecular Operating Environment – MOE program (ver 2018; Chemical Computing Group; Montreal, Canada). The molecules were energy minimized and geometry optimized for docking studies. Since carvones occupy a small volume and have the potential to bind nearly any binding pocket, we utilized a triage-based approach to finalize the predicted binding pocket. We screened the PAS-B domain of the AhR containing the binding pockets for TCDD, FICZ, BaP, CH-223191, Vemurafenib, Dabrafenib, PLX7904, PLX8394, Resveratrol as detailed in [18] and our newly developed methylindoles [25]. Pockets included TCDD as a control for each of these dockings. All docking screening experiments were performed using GOLD version 5.2 (Cambridge Crystallographic Data Centre, Cambridge, UK) [29]. The complexes were ranked using the default option of GOLD SCORE, and the best ranking complexes were visualized in MOE. The molecules were also docked to the AhR derived from the AhR-ARNT complex. S-carvone bound AhR was then energy minimized and subject to molecular dynamics simulation with a production run of 10ns. All simulations were performed using Amber99 forcefield as adopted in the MOE program.

### Microscale thermophoresis

A codon-optimized fragment of human AhR encoding amino acid residues 23–273 was synthesized and cloned into pET28b(+) using Ndel and BamHI restriction sites, expressing N-terminally fused 6×His-tag. A codon-optimized fragment of mouse ARNT encoding amino acid residues 85–345 was synthesized and cloned into pETDuet-1 using BamHI and HindIII restriction sites, expressing N-terminally fused 6×His-tag or using Ncol and HindIII restriction sites, expressing N-terminally FLAG-tag (GenScript, Leiden, Netherlands). A selection of truncated versions of the AhR and ARNT was done based on previously published data [27,30]. Both constructs were co-expressed in Rosetta 2 (DE3) *E. coli* cells (Novagen). Protein production was induced with 1 mM isopropyl-β-thiogalactopyranoside, and cells were grown at 20°C in LB medium overnight. Cells were lysed at 30 kpsi using the One-Shot cell lyser (Constant Systems Ltd.) and upon addition of EDTA-free cOmplete™ protease inhibitor cocktail (Roche). B-PER complete bacterial protein extraction reagent (Thermo) and Denerase (c-LEcta) were added to the lysate. Protein heterodimers were partially purified using HisPur Cobalt columns (Thermo Fisher Scientific) into the final buffer containing 20 mM HEPES, pH 7.0, 300 mM NaCl, and 5% (w/v) glycerol. The presence of AhR and ARNT proteins was verified by Western blot using anti-His-tag (mouse monoclonal, MA1-21315, dilution 1:1000, Invitrogen) anti-FLAG-tag (rabbit monoclonal, 14793S, dilution 1:1000, Cell Signaling Technology) antibodies. In parallel, lysates from E. coli were separated by electrophoresis using precast NuPAGE Bis-Tris protein gels (Thermo Fisher Scientific) and visualized by a Coomassie Brilliant Blue staining. Excised gel pieces with protein bands corresponding to the expected molecular masses of recombinant AhR and ARNT were processed using previous in-gel digestion and peptide extraction protocols [31].

Peptides from the digests were subjected to nanoflow liquid chromatography separations coupled to electrospray ionization tandem mass spectrometry (nanoLC-ESI-MS/MS) for protein identification *via* peptide sequencing and database searches. The nanoLC-ESI-MS/MS instrumental system was a Dionex UltiMate3000 RSLCnano liquid chromatography (Thermo Fisher Scientific) and an amaZon speed ETD ion trap equipped with a CaptiveSpray ion source (Bruker Daltonik). The chromatographic columns used and the mobile phases composition, gradient programming, data collection, and bioinformatics were already described [32].

Protein fractions were concentrated to 2 mg.ml^-1^ using 10 kDa filters (Amicon) and stored at 5°C for 10 days. Microscale thermophoresis was used to determine S-carvone and D-limonene’s binding to human 6×His-tagged hAhR in complex with FLAG-mARNT. The protein (200 nM) was fluorescently labeled using a RED-tris-NTA 2^nd^ generation dye (NanoTemper Technologies GmbH) and a 1:1 dye/protein molar ratio in the reaction buffer: 20 mM Tris-HCl, pH 7.4 supplemented with 150 mM NaCl and 0.075% Tween-20. Ligands were dissolved in ethanol (max. 0.5% final concentration in the reaction mixture). Measurements were performed on a Monolith NT.115 instrument (NanoTemper Technologies GmbH) at 25°C with 3 s / 22 s / 2 s laser off/on/off times and continuous sample fluorescence recording in premium capillaries and using an excitation power of 90% and a high MST power mode. The normalized fluorescence ΔF_norm_ [‰] as a function of the ligand concentration was analyzed to conclude a ligand binding interaction.

### Statistics

All statistical analyses, as well as the calculations of half-maximal effective concentration (EC_50_), EC_80_, and half-maximal inhibitory concentration (IC_50_) values, were performed using GraphPad Prism 8 for Windows (GraphPad Software, La Jolla, CA, U.S.A.). The number of independent repeats and technical replicates are stated in the respective figure legends for all the experiments. Where appropriate, data were processed by one-way analysis of variance (ANOVA) followed by Dunnett’s test. Results with p-values lower than 0.05 were considered significant—the EC_50_, EC_80_, and IC_50_ values were calculated using the nonlinear regression by the least-square fitting method. The R-squared value was checked in all of the calculations and did not drop below 0.9. Inhibition constant (K_i_) was calculated using the Cheng-Prusoff equation [33].

## RESULTS

### Carvones are non-competitive antagonists of the AhR

Human stably transfected reporter cell line AZ-AHR [23] was used to investigate the effects of carvones on transcriptional activity of the AhR. Model AhR full agonists, including TCDD, BaP, and FICZ, caused a concentration-dependent increase of AhR-mediated luciferase activity (Figure 1A). Carvones did not affect the basal transcriptional activity of AhR (Figure 1A). Effects of carvones on agonist-inducible AhR activity were examined in cells co-incubated with a fixed concentration of agonist ligands (corresponding to their EC_80_) and increasing concentration carvones (antagonist mode). Both carvones exerted concentration-dependent antagonist effects on AhR activation by all tested agonists. In the case of FICZ, the value of IC_50_ was not reached even in the highest concentrations of carvones. The inhibitor constants of S-carvone / R-carvone on BaP-and TCDD-inducible AhR activities were 14.0 μM /11.6 μM and 8.6 μM / 10.5 μM, respectively (Figure 1B). Next, we incubated cells with fixed concentrations of carvones (0 μM; 10 μM; 100 μM; 500 μM; 1000 μM) combined with increasing concentrations of AhR agonists. We observed a gradual decrease of E_MAX_ (and a slight decline of EC_50_) with increasing fixed concentrations of carvones for each tested agonist (Figure 1C). These data imply that carvones are primarily either non-competitive, irreversible competitive, or uncompetitive antagonists of the AhR. Since the same concentration of carvones antagonized both high and low concentrations of all used agonists to a comparable degree (Suppl. Table 1), the uncompetitive mechanism can be ruled out [34].

### Carvones down-regulate AhR target gene, *Cytochrome P450 1A1 (CYP1A1)*

Since carvones antagonized the AhR, we examined the effects of carvones on the ligand-inducible expression of prototypical AhR target gene CYP1A1, using a complementary set of AhR competent human cells, including hepatocarcinoma cells HepG2, colon adenocarcinoma cells LS180, and immortal keratinocytes HaCaT. Both carvones inhibited ligand-inducible expression of *CYP1A1* mRNA and protein in cell type-specific and ligand-selective manner. Induction of *CYP1A1* by TCDD was inhibited by carvones in all cell lines, by BaP in LS180 and HaCaT cells, and by FICZ only in HaCaT cells (Figure 2). We also observed down-regulation of TCDD-, BaP-and FICZ-inducible *CYP1A1* and *CYP1A2* mRNAs by carvones in three primary cultures of human hepatocytes (Suppl. Figure 1). Also, TCDD-inducible, the AhR receptor-regulated, 7-ethoxyresorufin-*O*-deethylase (EROD) was strongly decreased by S-carvone in AZ-AHR human hepatoma cells (Suppl. Figure 2).

**Figure 2.**
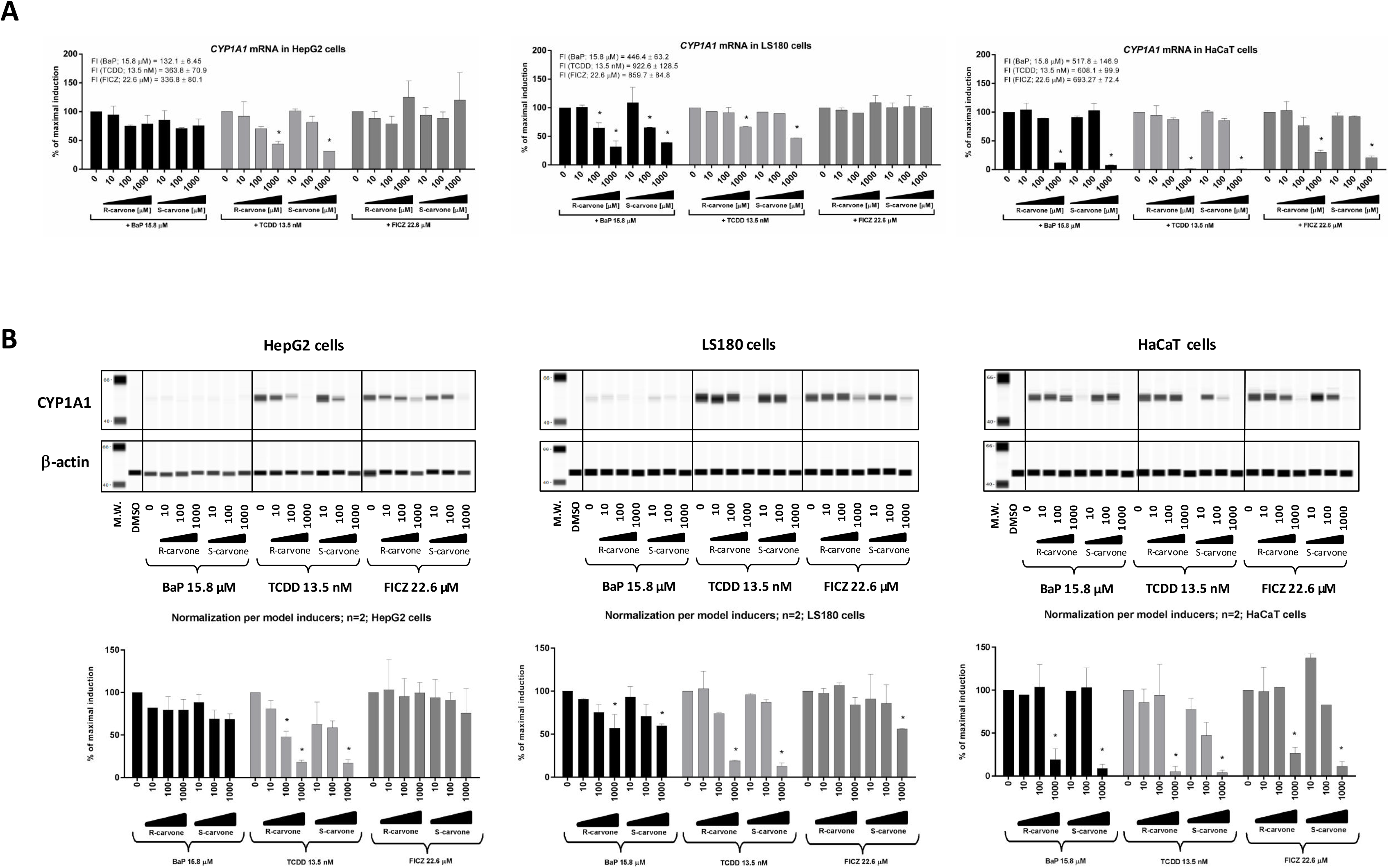
Down-regulation of CYP1A1 in human cell lines by carvones. HepG2, LS180, and HaCaT cells were incubated for 24 h with carvones (0 μM – 1000 μM) in the presence of AhR agonists TCDD, BaP and FICZ, applied in their EC_80_ concentrations. Incubations and measurements were performed in triplicates (technical replicates). **(A)** RT-PCR analyses of CYP1A1 mRNA; results expressed relative to agonist in the absence of carvones (100%). Data are mean ± S.D. from two independent cell passages. Results were normalized using GAPDH as a housekeeping gene. The absolute values of CYP1A1 mRNA fold inductions (F.I.) by model agonists are indicated in-text inserted in bar graphs from each cell line. **(B)** Quantitative automated western-blot analysis by SallySue of CYP1A1 protein. Representative SallySue records from one cell passage are shown. Bar graphs at the bottom show quantified CYP1A1 protein normalized *per* β-actin; data are expressed relative to agonist in the absence of carvones (100%) and are mean ± S.D. from two independent cell passages. * = significantly different from ligand in the absence of carvones (p<0.05)

### Carvones influence cellular functions of the AhR

We analyzed in detail the effects of carvones on individual cellular events throughout the AhR signaling pathway. The AhR agonists TCDD, BaP, and FICZ triggered the AhR from the cytosol to the nucleus, and carvones did not influence this process. Also, carvones alone did not induce AhR nuclear translocation (Figure 3A; Suppl. Table 2). Nuclear, ligand-bound AhR forms a heterodimer with ARNT, which in turn binds specific dioxin-response elements in promoters of target genes, such as *CYP1A1*. This pathway, involving ARNT, is referred to as canonical AhR signaling. Carvones strongly inhibited the formation of AhR-ARNT heterodimer (Figure 3B) and the binding of the AhR in *CYP1A1* promoter (Figure 3C) in cells stimulated with TCDD- and BaP-, but not with FICZ. A scheme summarizing the effects of carvone on the AhR functions is depicted in Figure 3D.

**Figure 3.**
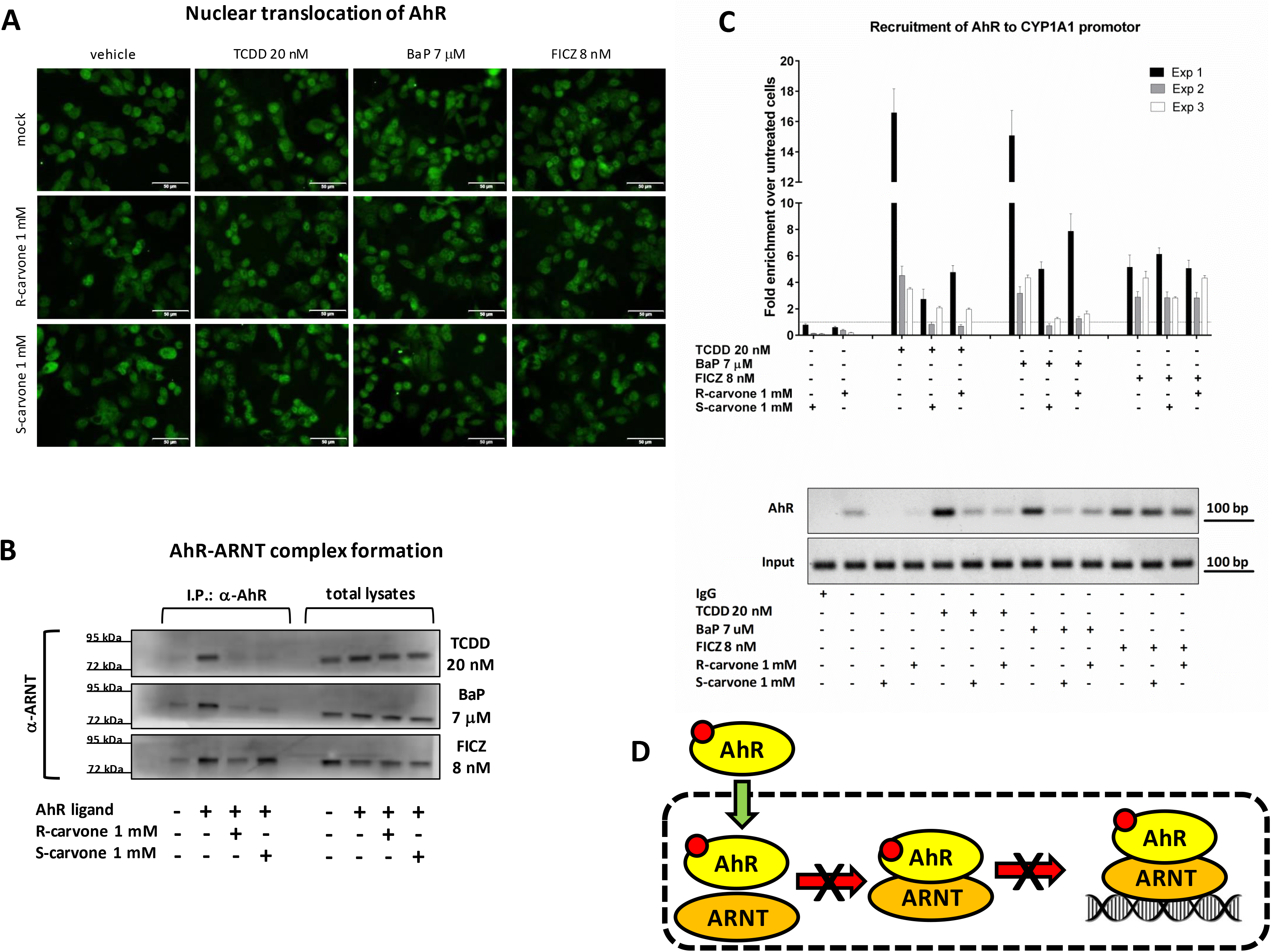
Carvones influence cellular functions of the AhR. Cells were incubated for 90 min with carvones (1000 μM) in combination with vehicle (0.1% DMSO) or AhR agonists TCDD (20 nM), BaP (7 μM), and FICZ (8 nM). **(A) Nuclear translocation of the AhR is not influenced by carvones.** Microscopic specimens from LS180 cells were prepared using Alexa Fluor 488 labeled primary antibody against AhR and DAPI. The AhR was visualized and evaluated using a fluorescence microscope. Experiments were performed in two consecutive cell passages, with all tested compounds in duplicates. The representative images are shown. **(B) Carvones inhibit the formation of AhR-ARNT heterodimer.** Protein co-immunoprecipitation – formation of AhR-ARNT heterodimer in LS180 cells. Representative immunoblots of immuno-precipitated protein eluates and total cell lysates are shown. Experiments were performed in three consecutive cell passages. **(C) Carvones inhibit the binding of the AhR in the CYP1A1 promoter.** Chromatin immunoprecipitation ChIP – binding of the AhR in CYP1A1 promoter in HepG2 cells. Bar graph (top) shows enrichment of CYP1A1 promotor with the AhR as compared to vehicle-treated cells. Representative DNA fragments amplified by PCR analyzed on a 2% agarose gel are from the 2^nd^ experiment (bottom). Experiments were performed in three consecutive cell passages. **(D) Schematic depiction of carvones’ cellular effects on the AhR**.

### S-carvone does not inhibit a random panel of protein kinases, including PKC

There are numerous reports about the involvement of protein kinase C in the AhR functions. Blocking protein kinase C (PKC) activity was reported to inhibit transcription of CYP1A1 but has no effect on nuclear translocation of the AhR [35], which was also the case here, observed with carvones. Therefore, we tested whether carvones inhibit PKC catalytic activity. We did not observe any decline in PKC activity measured in lysates from HepG2 cells incubated with carvones in concentrations up to 1000 μM, which rules out PKC inhibition’s involvement in the effects of carvones on the AhR (Figure 4A). We also evaluated the interaction between 100 μM S-carvone and 468 human protein kinases, employing KINOMEscan™ (scanMAX assay), a proprietary active site-directed competition binding assay [26]. The minimal inhibitory threshold by screening platform KINOMEscan™ is 35% of control kinase activity. Out of 468 kinases tested, 467 were above the 35% cut-off. TYK2(JH2 domain pseudo-kinase) activity was inhibited down-to 27% of control activity, but this kinase is not relevant to the regulation of the transcription activity (Figure 4B; Suppl. Figure 3). Overall, we excluded the possibility that the effects of carvones on the AhR signaling are indirect due to the inhibition of human kinome, particularly PKC.

**Figure 4.**
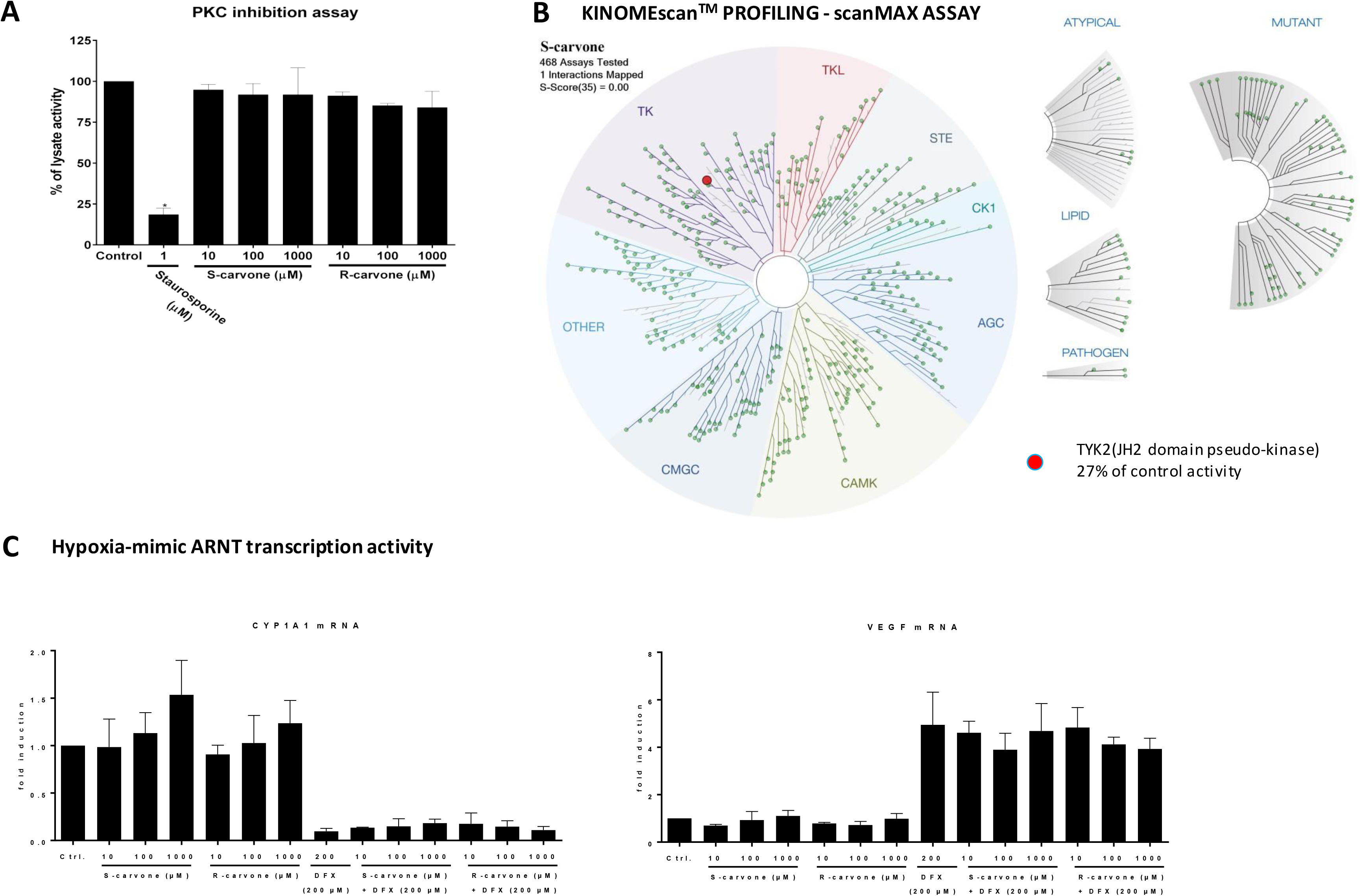
Evaluation of off-target effects of carvones. **(A) PKC inhibition assay**: Catalytic activity of PKC was measured in lysates from HepG2 cells incubated with vehicle (DMSO, 0.1% V/V), staurosporine (1μM), and carvones (10 μM; 100 μM; 1000 μM). Data are mean ± S.D. from three independent experiments. Incubations and measurements were performed in uniplicates (technical replicates). **(B) KINOMEscan™ profiling**: The interaction between 100 μM S-carvone and 468 human protein kinases, employing KINOMEscan™ (scanMAX assay), proprietary active site-directed competition binding assay. A low-resolution interaction map is shown. **(C) Hypoxia-mimic VEGF induction**: HaCaT cells were incubated for 24 h with carvones (10 μM; 100 μM; 1000 μM) in combination with vehicle (0.1% DMSO) or deferoxamine (DFX; 200 μM). The expression of VEGF and CYP1A1 mRNAs was measured using RT-PCR. Incubations and measurements were performed in duplicates (technical replicates). Data are mean ± S.D. from three independent cell passages and are expressed as fold induction over the vehicle-treated cells. Results were normalized using GAPDH as a housekeeping gene.

### Carvones do not inhibit the transcriptional activity of ARNT

ARNT is involved in other cellular pathways besides that of AhR, such as hypoxia signaling that is transcriptionally mediated by ARNT heterodimer with hypoxia-inducible factor 1α (HIF1α). Therefore, we investigated the effects of carvones on hypoxia-mimic inducible, ARNT-dependent expression of vascular endothelial growth factor (VEGF) mRNA in HaCaT cells incubated with deferoxamine. The levels of VEGF mRNA were induced 5-fold by deferoxamine, and carvones did not influence this induction in concentrations up to 1000 μM (Figure 4C-right panel). Consistently, the hypoxia-mimic decrease of CYP1A1 mRNA was not affected by carvones (Figure 4C-left panel). These data imply that carvones do not inhibit ARNT transcriptional activity and that disruption of AhR-ARNT complex formation is not due to the interaction of carvones with ARNT.

### Binding of carvones to the AhR

Reporter gene assay revealed that carvones are non-competitive antagonists of the AhR (Figure 1), implying they should not competitively displace ligands from binding at the AhR. This assumption was corroborated by competitive radio-ligand binding assay, where S-carvone did not inhibit the binding of ^3^H-TCDD at mouse hepatic AhR. However, we observed a slight, concentration-independent decrease of ^3^H-TCDD binding in the presence of 1000 μM S-carvone (Figure 5A). Non-competitive antagonism may occur through (i) Allosteric hindrance (direct or involving conformational change) of ligand binding pocket at AhR, thereby preventing proper binding of the ligand and switching-on the AhR. This scenario is unlikely because the ligand-dependent nuclear translocation of AhR was not disturbed by carvones (Figure 3A). For this reason, also irreversible competitive antagonism is not likely. (ii) An indirect mechanism, occurring either at AhR or off-target, such as protein kinases or ARNT (*vide supra*). Therefore, we further investigated the allosteric binding of carvones at the AhR and their effects on AhR-ARNT heterodimerization. Molecular Docking of carvones to various known binding pockets of the AhR ligands such as TCDD, resveratrol, FICZ, BaP, and methylindoles suggested that carvones may non-specifically bind to these sites with an average docking score of 47.5 and 42, respectively. However, this binding could be due to their relatively small size and could have no functional effect. Based on the experimental evidence that carvones inhibit the formation of AhR-ARNT (Figure 3B), we docked these molecules to the heterodimerization interface of AhR and ARNT. This interface spans several interdomain interactions that also form the dioxin responsive element binding pocket [36]. One such interface region is the α1-α2 helical region of the bHLH domain consisting of residues Leu43, Leu47, Leu50 from α1 helix and Tyr76, Leu72, Leu77 from α2 helical region. Carvones were docked to the interface site, and the complex of the AhR with carvones was simulated for 10 ns to allow the ligand to stably dock to the AhR (Figure 5B; left). Carvones bind favorably at a site formed by residues from the bHLH domain, including close contact with Tyr76, Pro55, Phe83, Tyr137, Leu72, Ala138, Lys80, Ser75, Phe56, Ala79, Phe136, Gln150 and Ile154 (Figure 5B; middle). More significantly, binding of carvones to this site shifts the position of both the α1 and α2 helical region by 1-3 Å (Figure 5B; right) that can significantly affect the formation of the AhR-ARNT complex. Using microscale thermophoresis, using bacterially co-expressed fragments of the AhR and ARNT, we showed that carvones bind the AhR but not ARNT (Figure 5C). Whereas the binding of carvones to the AhR was concentration-dependent, the apparent binding constant K_D_ could not be determined since it lay in the low millimolar range, probably due to artificial conditions using truncated variants of the AhR and ARNT. The AhR fragment spanned from 23 to 273 amino acid residues, which implies that the binding of carvones was localized outside the conventional ligand-binding domain, but within the bHLH/PAS-A region of the AhR. These data fully support the hypothesis that carvones’ non-competitive antagonism involves their allosteric binding at the AhR. Also, D-limonene (de-oxo analog of carvone) did not display the AhR antagonism and did not bind the AhR, which reveals the significance of *oxo* moiety in carvone molecule for its interaction with the AhR, tentatively through hydrogen bonds (Suppl. Figure 4).

**Figure 5.**
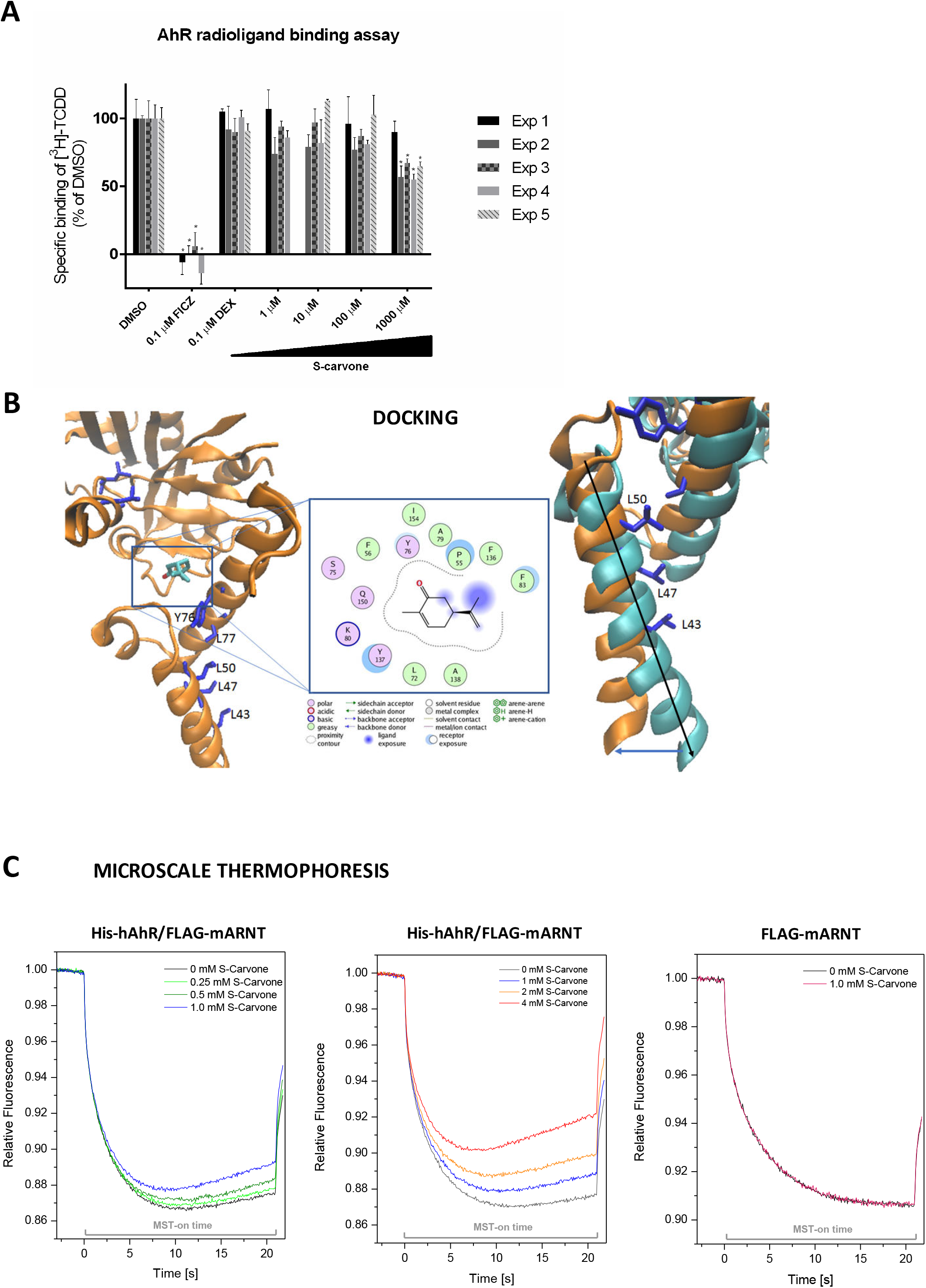
Binding of S-carvone at the AhR. **(A) Competitive radioligand binding assay**: Cytosolic protein (2 mg/mL) from Hepa1c1c7 cells was incubated with S-carvone (1 μM, 10 μM, 100 μM, 1000 μM), FICZ (10 nM), DEX (100 nM; negative control) or DMSO (0.1% V/V; corresponds to *specific binding of [^3^H]-TCDD = 100%*) in the presence of 2 nM [^3^H]-TCDD. Specific binding of [^3^H]-TCDD was determined as a difference between total and non-specific (200 nM; 2,3,7,8-tetrachlorodibenzofuran) reactions. The significance (p<0.05) was tested against negative control (*). Five independent experiments were performed, and the incubations and measurements were done in triplicates in each experiment (technical replicates). The error bars represent the mean ± S.D. **(B) Molecular docking**: S-carvone (licorice stick and colored atom type, Carbon = cyan and Oxygen = red) binds to a site proximal to the heterodimerization interface of AhR (depicted as orange ribbons with interface residues shown as blue licorice sticks and labeled; left panel) with residues from α1 and α2 helices contributing to the binding interactions (center panel). Binding of S-Carvone to this site also leads to conformational changes in α1 and α2 helices (new positions shown as cyan ribbons; right panel), thereby disrupting the formation of AhR-ARNT interface**. (C) Microscale thermophoresis**; *left panel:* co-expressed hAhR-His + mARNT-Flag incubated with vehicle and/or S-carvone (0.25 mM, 0.5 mM, 1 mM); *central panel:* co-expressed His-hAhR + FLAG-mARNT incubated with vehicle and/or S-carvone (1 mM, 2 mM, 4 mM); *right panel:*. FLAG-mARNT incubated with the vehicle or 1 mM S-carvone.

## DISCUSSION

Therapeutic targeting of the AhR has long been neglected, mainly due to the negative stigma of being a receptor mediating TCDD toxicity. With increasing knowledge on the physiological and pathophysiological roles of the AhR, the attempts for its targeting have emerged, including the therapy of cancer, inflammatory bowel disease, or atopic dermatitis. Following strategies are employed: (i) A repositioning of clinically used AhR-active drugs (e.g., tranilast, flutamide, omeprazole); (ii). Chemoprevention with dietary AhR-active compounds (e.g., indole-3-carbinol, diindolylmethane); (iii) Application of novel AhR ligands identified by screening chemical libraries (e.g., CH223191) or by rational design (e.g., PY109).

The interactions between the small-molecule compound and the AhR may occur either directly (ligand-dependent) or indirectly (ligand-independent) through off-targets such as PKC [35], protein tyrosine kinases [37], or cAMP [38]. To date, all reported AhR ligands, both agonists and antagonists, are the orthosteric ones. i.e., those that bind to a conventional discrete site on the AhR protein are referred to as ligand-binding pocket. Recently, three structurally distinct groups of chemicals were defined according to the mode of their interactions with residues within the AhR ligand binding site [39]. Depending on their effects on the AhR functions, these ligands comprise full agonists, partial agonists, and competitive antagonists. Herein, this is the first report on small molecule compounds acting as allosteric antagonists of human AhR, which may potentially be of clinical importance. Indeed, targeting secondary binding sites at a variety of receptors is an emerging approach in drug discovery [40–42]. The examples of already approved allosteric modulator drugs are Cinacalcet for the treatment of hyperparathyroidism (positive modulator of the calcium-sensing receptor) or Maraviroc for the treatments of AIDS (negative modulator of C-C chemokine receptor 5). Many other compounds are yet under patent protection, such as positive allosteric modulators of dopamine receptors in the treatment of Parkinson’s disease and schizophrenia (Pat. Appl. # WO/2014/193781; Eli Lilly&Co) or negative allosteric modulators of metabotropic glutamate receptors for the treatment of CNS disorders (Pat. Appl. # WO/2014/195311; Janssen Pharmaceutica). Scheuermann et al described synthetic allosteric inhibitors of hypoxia-inducible factor HIF2α, which bound in a large cavity within a hydrophobic core of PAS-B domain of HIF2α, inducing structural and functional changes leading to the antagonization of HIF2α heterodimerization with ARNT [43].

Applying a series of complementary mechanistic experiments, we demonstrate that carvones are non-competitive antagonists of human AhR, acting through allosteric binding in the region of the AhR involved in heterodimerization with ARNT, thereby preventing the formation of functional AhR-ARNT heterodimer. In brief, detailed analyses of the AhR transcriptional response in reporter gene assays revealed a non-competitive mechanism of carvones antagonism. This is consistent with the finding that carvones did not displace ^3^H-TCDD from binding at AhR and also did not inhibit ligand-elicited nuclear translocation of the AhR. On the other hand, S/R-carvones inhibited the formation of AhR-ARNT heterodimer, and all downward events involving binding of the AhR to DNA, and the expression of AhR-target genes. In search of the exact mechanism of how carvones inhibit the formation of AhR-ARNT heterodimer, we excluded the interaction of carvones with potential off-targets, including ARNT, PKC, and other 468 kinases.

Differential roles of the AhR and ARNT residues on molecular events preceding (ligand binding, nuclear translocation) and following (DNA binding) heterodimerization of the AhR with ARNT were determined by Corrada et al. by combining site-directed mutagenesis, structural modeling, and homology docking approach [44,45]. Crystal structure of mouse AhR PAS-A domain revealed that mouse AhR residues Ala119 and Leu120 are critically crucial for hydrophobic interactions at the AhR-ARNT interface and the process dimerization [46]. Seok et al. determined the crystal structure of mouse AhR-ARNT heterodimer in complex with DRE, showing that ARNT curls around AhR into a highly inter-winded asymmetric architecture, with extensive heterodimerization interfaces and AhR inter-domain interactions. They proposed the phenomenon of ligand-selective structural hierarchy for complex scenarios of the AhR activation [36]. Mutations in mAhR residues Leu42 and Leu120 (homologous to human Leu43 and Leu122) led to a decreased binding of AhR-ARNT to DRE [36], which corroborates the findings of Wu et al. [46]. Interestingly, mutation of Leu49 in mAhR kept intact nuclear translocation of the AhR but inhibited its transcription activity [36], which is mimicked by binding of carvones at the AhR. According to our docking data, carvones interact with residues from the bHLH domain, including close contact with Tyr76, Pro55, Phe83, Tyr137, Leu72, Ala138, Lys80, Ser75, Phe56, Ala79, Phe136, Gln150, and Ile154. Binding of carvones to this site shifts the position of α1 and α2 helical regions by 1-3 Å (Figure 5B; right) that can significantly affect the formation of the AhR-ARNT complex. This assumption was experimentally confirmed, and direct binding of carvones at the AhR fragment spanning from 23 to 273 amino acid residues was demonstrated using microscale thermophoresis. Also, the significance of *oxo* moiety in the molecule of carvone, for its interaction with the AhR, tentatively through hydrogen bonds, was demonstrated.

Two issues about the AhR-active concentration of carvones: Firstly, the biological effects of carvones against the AhR were attained in concentrations spanning from 100 μM to 1000 μM, which might appear high; however, the available data suggest that these concentrations are relevant. Topical application of 300 mg of R-carvone or S-carvone, which are used as skin permeabilizers in transdermal patches, resulted in maximal plasma concentrations of ~0.6 μM and ~0.2 μM, respectively [22]. On the other hand, local concentrations of carvones in keratinocytes, following the topical application, must be in orders of magnitude higher than plasma levels. Blood levels of carvone in volunteers, who received 100 mg of caraway oil (~54.5 mg carvone) in coated capsules, reached approx. 0.1 μM [47]. However, local concentrations of carvones in enterocytes (intestinal first pass) and hepatocytes (hepatic first-pass) must be much higher than those reached in plasma. The concentration of carvones in meals is approx. 150 μM, implying exposure of enterocytes to those concentrations when consuming food containing EOs of caraway, spearmint, or dill [48]. Also, European Food Safety Authority EFSA defined acceptable daily intake of S-carvone as 0.6 mg/kg of body weight. Besides, a recent estimate based on recommended dose and published a fecal excreted fraction of 200 marketed drugs report a median expected colon concentration of 80 μM for drugs having median serum concentration of 0.6 μM, implying globally >100-times higher drug concentrations in the gut as compared to blood [49]. Collectively, potential clinical or preventive use of carvones as the AhR antagonists is predestined by their local effects (not systemic ones) on the skin (topical application) or in the intestines (peroral intake). Secondly, carvones’ antagonism at the level of gene expression (mRNA, protein, EROD, luciferase) occurred with IC_50_ values approx. From 10^-5^ M to 10^-4^ M, whereas 10^-3^ M carvones antagonized the AhR functions (heterodimerization, DNA binding). The plausible explanation for this discrepancy could be differential cellular uptake of carvones, when the gene expression and the AhR functions were studied after 24 h and 90 min of incubation, respectively.

In summary, we report here dietary monocyclic monoterpenoid carvones, as a new class of non-competitive antagonists of the AhR, acting through allosteric binding at the AhR, thereby blocking heterodimerization with ARNT and constraining transcriptional functions of AhR-ARNT. While hundreds of orthosteric AhR ligands, including antagonists, were described, this is the first report on allosteric antagonism of the AhR by small-molecule compounds, which might be of clinical but also fundamental mechanistic importance.

## Abbreviations

AhR: Aryl Hydrocarbon Receptor
ARNT: AhR Nuclear Translocator
BaP: Benzo[a]pyrene
DEX: Dexamethasone
EROD: 7-ethoxyresorufin-*O*-deethylase
FICZ: 6-formylindolo[3,2-b]carbazole
OR1A1: Odorant Receptor 1A1
PKC: Protein Kinase C
TCDD: 2,3,7,8-tetrachlorodibenzo-*p*-dioxin
TCDF: 2,3,7,8-tetrachlorodibenzofuran
VEGF: Vascular Endothelial Growth Factor

## ACKNOWLEDGEMENTS

Financial support from Czech Health Research Council [NV19-05-00220], the student grant from Palacky University in Olomouc [PrF-2020-006], Juergen Manchot Foundation (to K.M.R.), NIH-R01CA222469, Department of Defense Partnering PI (W81XWH-17-1-0479; PR160167), (ES030197) (to S.M.) is acknowledged. We thank Dr. Radka Končitíková for the assistance with microscale thermophoresis experiments.

## AUTHORS CONTRIBUTIONS

*Participated in research design:* Z.D., T.H.S., S.M.

*Conducted experiments:* K.P., B.V., I.Z., K.K., E.J., R.V., K.M.R., S.K., D.K., M.K., M.Š.

*Contributed new reagents and analytic tools:* Z.D., T.H.S., S.K.

*Performed data analysis:* K.P., B.V., I.Z., K.K., E.J., R.V., J.V., K.M.R., S.K., Z.D., D.K., M.K., M.Š.

*Wrote or contributed to the writing of the manuscript:* Z.D., T.H.S., S.M., S.K.

## DATA AVAILABILITY

All data needed to evaluate the paper’s conclusion are present in the paper or the Supplementary Materials. Additional datasets generated during or analyzed during the current study are available from the corresponding author on reasonable request.

## MATERIALS & CORRESPONDENCE

Zdeněk Dvořák

Department of Cell Biology and Genetics Faculty of Science, Palacky University Olomouc Slechtitelu 27; 783 71 Olomouc; Czech Republic

E: T: +420-58-5634903 F: +420-58-5634901

Sridhar Mani

Department of Genetics and Department of Medicine Albert Einstein College of Medicine

Bronx, NY 10461, U.S.A.

E

---

**Supplementary Figure 1. Down-regulation of CYP1A1 and CYP1A2 mRNAs in primary cultures of human hepatocytes.** Human hepatocytes cultures from three tissue donors (LH75, HEP200571, HEP220993) were incubated for 24 h with carvones (10 μM, 100 μM, 1000 μM) in the presence of AhR agonists TCDD (5 nM, 50 nM), BaP (1 μM, 10 μM) and FICZ (10 nM, 100 nM). RT-PCR quantified CYP1As mRNAs. **(A)** Fold induction of CYP1A genes by AhR agonists; **(B)** Percentage of CYP1As maximal induction by model agonists in the presence of carvones. Data were normalized using GAPDH as a housekeeping gene.

**Supplementary Figure 2. Down-regulation of 7-ethoxyresorufin-*O*-deethylase EROD by S-carvone in hepatoma cells.** Catalytic activity of EROD was measured using fluorescent substrate in AZ-AHR cells pre-incubated for 24 h with vehicle (DMSO; 0.1% v/v), TCDD (13.5 nM) and/or mixture of S-carvone (1 mM) + TCDD (13.5 nM). Incubations and measurements were performed in triplicates (technical replicates). The bar graph data are mean ± S.D. from three consecutive cell passages and are expressed as the percentage of fluorescence in TCDD-treated cells.

**Supplementary Figure 3. KINOMEscan™ profiling:** The interaction between 100 μM S-carvone and 468 human protein kinases, employing KINOMEscan™ (scanMAX assay), proprietary active site-directed competition binding assay. A high-resolution interaction map is shown.

**Supplementary Figure 4. D-limonene does not interact with the AhR. (A) Chemical structures** of S-carvone and D-limonene and their schematic interaction with the AhR; **(B) Reporter gene assay** was carried out in AZ-AHR cells, incubated for 24 h with increasing concentrations of D-limonene in combination with the AhR agonists TCDD (13.5 nM), BaP (15.8 μM) and FICZ (22.6 μM). Incubations and measurements were performed in quadruplicates (technical replicates). The bar graph shows the percentage of maximal induction attained by a model AhR agonist. Data are mean ± S.D. from three consecutive cell passages. * = significantly different from AhR agonist in the absence of D-limonene (p<0.05); **(C) Microscale thermophoresis** using co-expressed His-hAhR + FLAG-mARNT incubated with vehicle (upper panel) or 1 mM D-limonene (lower panel).

**Supplementary Table 1. Quantitative analysis of S/R-carvones in reporter gene assay against uncompetitive antagonism.** Reporter gene assay was carried out in stably transfected AZ-AHR cells, incubated for 24 h with a fixed concentration of S/R-carvones combined with increasing concentrations of AhR agonists. Experiments were performed in two independent cell passages. Incubations and measurements were performed in quadruplicates (technical replicates). Percentage of inhibition by S/R-carvones (10 μM; 100 μM; 500 μM; 1000 μM) was calculated for all tested agonists as follows: % Inhibition C_x_ = 100*(CARVONE C_0_ - CARVONE C_x_)/CARVONE C_0_

**Supplementary Table 2. Carvones do not influence the nuclear translocation of AhR.** LS180 cells were incubated for 90 min with S/R-carvones (1000 mM) in combination with vehicle (0.1% DMSO) or AhR agonists TCDD (20 nM), BaP (7 μM), and FICZ (8 nM). Microscopic specimens were prepared using Alexa Fluor 488 labeled primary antibody against AhR and DAPI. AhR was visualized and evaluated using a fluorescence microscope. Experiments were performed in two consecutive cell passages, with all tested compounds in duplicates. The table shows the total and AhR-positive counts of cells.

## REFERENCES

1. Stejskalova L, Dvorak Z, Pavek P (2011) Endogenous and exogenous ligands of aryl hydrocarbon receptor: current state of art. Curr Drug Metab 12: 198–212.

2. Angelos MG, Kaufman DS (2018) Advances in the role of the aryl hydrocarbon receptor to regulate early hematopoietic development. Curr Opin Hematol 25: 273–278.

3. Bock KW (2018) From TCDD-mediated toxicity to searches of physiologic AHR functions. Biochem Pharmacol 155: 419–424.

4. Gutierrez-Vazquez C, Quintana FJ (2018) Regulation of the Immune Response by the Aryl Hydrocarbon Receptor. Immunity 48: 19–33.

5. Fang ZZ, Krausz KW, Nagaoka K, Tanaka N, Gowda K, et al. (2014) In vivo effects of the pure aryl hydrocarbon receptor antagonist GNF-351 after oral administration are limited to the gastrointestinal tract. Br J Pharmacol 171: 1735–1746.

6. Ghotbaddini M, Moultrie V, Powell JB (2018) Constitutive Aryl Hydrocarbon Receptor Signaling in Prostate Cancer Progression. J Cancer Treatment Diagn 2: 11–16.

7. Chen J, Haller CA, Jernigan FE, Koerner SK, Wong DJ, et al. (2020) Modulation of lymphocyte-mediated tissue repair by rational design of heterocyclic aryl hydrocarbon receptor agonists. Sci Adv 6: eaay8230.

8. Safe S, Cheng Y, Jin UH (2017) The Aryl Hydrocarbon Receptor (AhR) as a Drug Target for Cancer Chemotherapy. Curr Opin Toxicol 2: 24–29.

9. Wilhelm SM, Rjater RG, Kale-Pradhan PB (2013) Perils and pitfalls of long-term effects of proton pump inhibitors. Expert Rev Clin Pharmacol 6: 443–451.

10. Dvorak Z, Vrzal R, Henklova P, Jancova P, Anzenbacherova E, et al. (2008) JNK inhibitor SP600125 is a partial agonist of human aryl hydrocarbon receptor and induces CYP1A1 and CYP1A2 genes in primary human hepatocytes. Biochem Pharmacol 75: 580–588.

11. Lu YF, Santostefano M, Cunningham BD, Threadgill MD, Safe S (1995) Identification of 3’-methoxy-4’-nitroflavone as a pure aryl hydrocarbon (Ah) receptor antagonist and evidence for more than one form of the nuclear Ah receptor in MCF-7 human breast cancer cells. Arch Biochem Biophys 316: 470–477.

12. Zhou J, Gasiewicz TA (2003) 3‘-methoxy-4’-nitroflavone, a reported aryl hydrocarbon receptor antagonist, enhances Cyp1a1 transcription by a dioxin responsive element-dependent mechanism. Arch Biochem Biophys 416: 68–80.

13. Kim SH, Henry EC, Kim DK, Kim YH, Shin KJ, et al. (2006) Novel compound 2-methyl-2H-pyrazole-3-carboxylic acid (2-methyl-4-o-tolylazo-phenyl)-amide (CH-223191) prevents 2,3,7,8-TCDD-induced toxicity by antagonizing the aryl hydrocarbon receptor. Mol Pharmacol 69: 1871–1878.

14. Choi EY, Lee H, Dingle RW, Kim KB, Swanson HI (2012) Development of novel CH223191-based antagonists of the aryl hydrocarbon receptor. Mol Pharmacol 81: 3–11.

15. Zhao B, Degroot DE, Hayashi A, He G, Denison MS (2010) CH223191 is a ligand-selective antagonist of the Ah (Dioxin) receptor. Toxicol Sci 117: 393–403.

16. Smith KJ, Murray IA, Tanos R, Tellew J, Boitano AE, et al. (2011) Identification of a high-affinity ligand that exhibits complete aryl hydrocarbon receptor antagonism. J Pharmacol Exp Ther 338: 318–327.

17. Bianchi-Smiraglia A, Bagati A, Fink EE, Affronti HC, Lipchick BC, et al. (2018) Inhibition of the aryl hydrocarbon receptor/polyamine biosynthesis axis suppresses multiple myeloma. J Clin Invest 128: 4682–4696.

18. Corre S, Tardif N, Mouchet N, Leclair HM, Boussemart L, et al. (2018) Sustained activation of the Aryl hydrocarbon Receptor transcription factor promotes resistance to BRAF-inhibitors in melanoma. Nat Commun 9: 4775.

19. Hawerkamp HC, Kislat A, Gerber PA, Pollet M, Rolfes KM, et al. (2019) Vemurafenib acts as an aryl hydrocarbon receptor antagonist: Implications for inflammatory cutaneous adverse events. Allergy 74: 2437–2448.

20. Bartonkova I, Dvorak Z (2018) Essential oils of culinary herbs and spices display agonist and antagonist activities at human aryl hydrocarbon receptor AhR. Food Chem Toxicol 111: 374–384.

21. Geithe C, Protze J, Kreuchwig F, Krause G, Krautwurst D (2017) Structural determinants of a conserved enantiomer-selective carvone binding pocket in the human odorant receptor OR1A1. Cell Mol Life Sci 74: 4209–4229.

22. Jager W, Mayer M, Reznicek G, Buchbauer G (2001) Percutaneous absorption of the montoterperne carvone: implication of stereoselective metabolism on blood levels. J Pharm Pharmacol 53: 637–642.

23. Novotna A, Pavek P, Dvorak Z (2011) Novel stably transfected gene reporter human hepatoma cell line for assessment of aryl hydrocarbon receptor transcriptional activity: construction and characterization. Environ Sci Technol 45: 10133–10139.

24. Denison MS, Rogers JM, Rushing SR, Jones CL, Tetangco SC, et al. (2002) Analysis of the aryl hydrocarbon receptor (AhR) signal transduction pathway. Curr Protoc Toxicol Chapter 4: Unit4 8.

25. Stepankova M, Bartonkova I, Jiskrova E, Vrzal R, Mani S, et al. (2018) Methylindoles and Methoxyindoles are Agonists and Antagonists of Human Aryl Hydrocarbon Receptor. Mol Pharmacol 93: 631–644.

26. Fabian MA, Biggs WH3rd, Treiber DK, Atteridge CE, Azimioara MD, et al. (2005) A small molecule-kinase interaction map for clinical kinase inhibitors. Nat Biotechnol 23: 329–336.

27. Schulte KW, Green E, Wilz A, Platten M, Daumke O (2017) Structural Basis for Aryl Hydrocarbon Receptor-Mediated Gene Activation. Structure 25: 1025–1033 e1023.

28. Wu D, Su X, Potluri N, Kim Y, Rastinejad F (2016) NPAS1-ARNT and NPAS3-ARNT crystal structures implicate the bHLH-PAS family as multi-ligand binding transcription factors. Elife 5.

29. Jones G, Willett P, Glen RC (1995) Molecular recognition of receptor sites using a genetic algorithm with a description of desolvation. J Mol Biol 245: 43–53.

30. Puyskens A, Stinn A, van der Vaart M, Kreuchwig A, Protze J, et al. (2020) Aryl Hydrocarbon Receptor Modulation by Tuberculosis Drugs Impairs Host Defense and Treatment Outcomes. Cell Host Microbe 27: 238–248 e237.

31. Shevchenko A, Tomas H, Havlis J, Olsen JV, Mann M (2006) In-gel digestion for mass spectrometric characterization of proteins and proteomes. Nat Protoc 1: 2856–2860.

32. Perutka Z, Sufeisl M, Strnad M, Sebela M (2019) High-proline proteins in experimental hazy white wine produced from partially botrytized grapes. Biotechnol Appl Biochem 66: 398–411.

33. Cheng Y, Prusoff WH (1973) Relationship between the inhibition constant (K1) and the concentration of inhibitor which causes 50 per cent inhibition (I50) of an enzymatic reaction. Biochem Pharmacol 22: 3099–3108.

34. Chen HSV, Pellegrini JW, Aggarwal SK, Lei SZ, Warach S, et al. (1992) Open-Channel Block of N-Methyl-D-Aspartate (Nmda) Responses by Memantine-Therapeutic Advantage against Nmda Receptor-Mediated Neurotoxicity. Journal of Neuroscience 12: 4427–4436.

35. Long WP, Pray-Grant M, Tsai JC, Perdew GH (1998) Protein kinase C activity is required for aryl hydrocarbon receptor pathway-mediated signal transduction. Mol Pharmacol 53: 691–700.

36. Seok SH, Lee W, Jiang L, Molugu K, Zheng A, et al. (2017) Structural hierarchy controlling dimerization and target DNA recognition in the AHR transcriptional complex. Proc Natl Acad Sci U S A 114: 5431–5436.

37. Backlund M, Ingelman-Sundberg M (2005) Regulation of aryl hydrocarbon receptor signal transduction by protein tyrosine kinases. Cell Signal 17: 39–48.

38. Oesch-Bartlomowicz B, Huelster A, Wiss O, Antoniou-Lipfert P, Dietrich C, et al. (2005) Aryl hydrocarbon receptor activation by cAMP vs. dioxin: divergent signaling pathways. Proc Natl Acad Sci U S A 102: 9218–9223.

39. Giani Tagliabue S, Faber SC, Motta S, Denison MS, Bonati L (2019) Modeling the binding of diverse ligands within the Ah receptor ligand binding domain. Sci Rep 9: 10693.

40. Abdel-Magid AF (2015) Allosteric modulators: an emerging concept in drug discovery. ACS Med Chem Lett 6: 104–107.

41. Venkatesh M, Wang H, Cayer J, Leroux M, Salvail D, et al. (2011) In vivo and in vitro characterization of a first-in-class novel azole analog that targets pregnane X receptor activation. Mol Pharmacol 80: 124–135.

42. Ekins S, Chang C, Mani S, Krasowski MD, Reschly EJ, et al. (2007) Human pregnane X receptor antagonists and agonists define molecular requirements for different binding sites. Mol Pharmacol 72: 592–603.

43. Scheuermann TH, Li Q, Ma HW, Key J, Zhang L, et al. (2013) Allosteric inhibition of hypoxia inducible factor-2 with small molecules. Nat Chem Biol 9: 271–276.

44. Corrada D, Soshilov AA, Denison MS, Bonati L (2016) Deciphering Dimerization Modes of PAS Domains: Computational and Experimental Analyses of the AhR:ARNT Complex Reveal New Insights Into the Mechanisms of AhR Transformation. PLoS Comput Biol 12: e1004981.

45. Corrada D, Denison MS, Bonati L (2017) Structural modeling of the AhR:ARNT complex in the bHLH-PASA-PASB region elucidates the key determinants of dimerization. Mol Biosyst 13: 981–990.

46. Wu D, Potluri N, Kim Y, Rastinejad F (2013) Structure and dimerization properties of the aryl hydrocarbon receptor PAS-A domain. Mol Cell Biol 33: 4346–4356.

47. Mascher H, Kikuta C, Schiel H (2001) Pharmacokinetics of menthol and carvone after administration of an enteric coated formulation containing peppermint oil and caraway oil. Arzneimittelforschung 51: 465–469.

48. Bartonkova I, Dvorak Z (2018) Essential oils of culinary herbs and spices activate PXR and induce CYP3A4 in human intestinal and hepatic in vitro models. Toxicol Lett 296: 1–9.

49. Maier L, Pruteanu M, Kuhn M, Zeller G, Telzerow A, et al. (2018) Extensive impact of non-antibiotic drugs on human gut bacteria. Nature 555: 623–628.

