## Supplementary data for "Dietary Monoterpenoids As a New Class of Allosteric Human Aryl Hydrocarbon Receptor Antagonists"

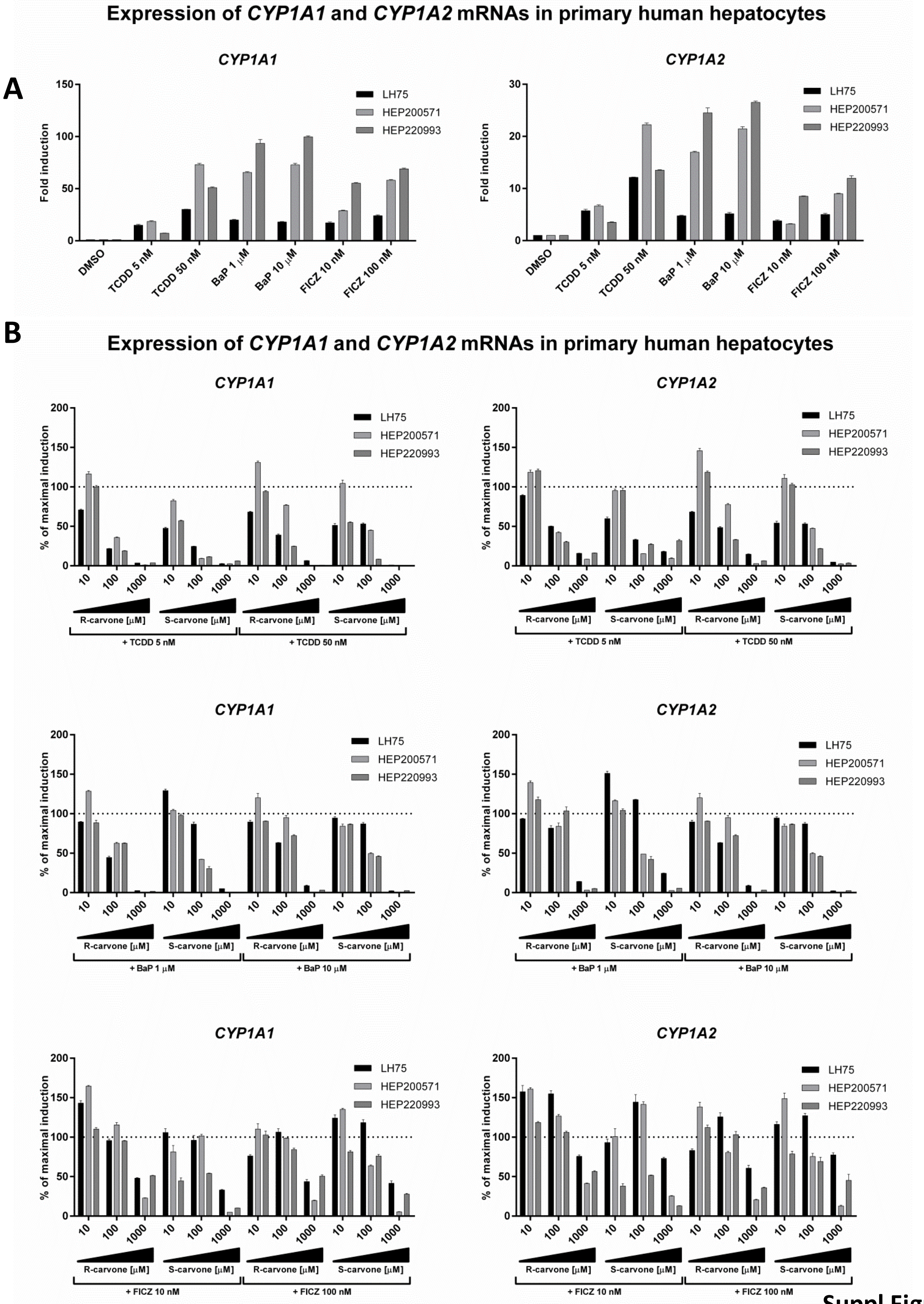


Suppl Figure 1

Supplementary Figure 2

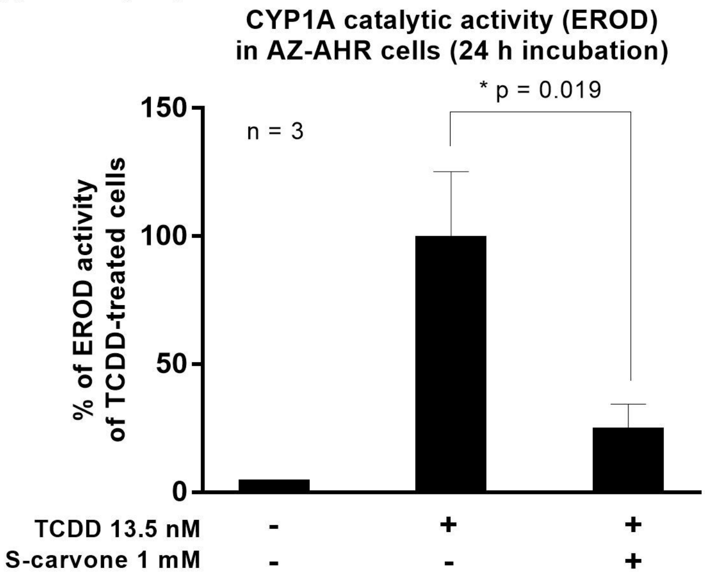

### S-carvone

468 Assays Tested
1 Interactions Mapped
S-Score(35) = 0.00

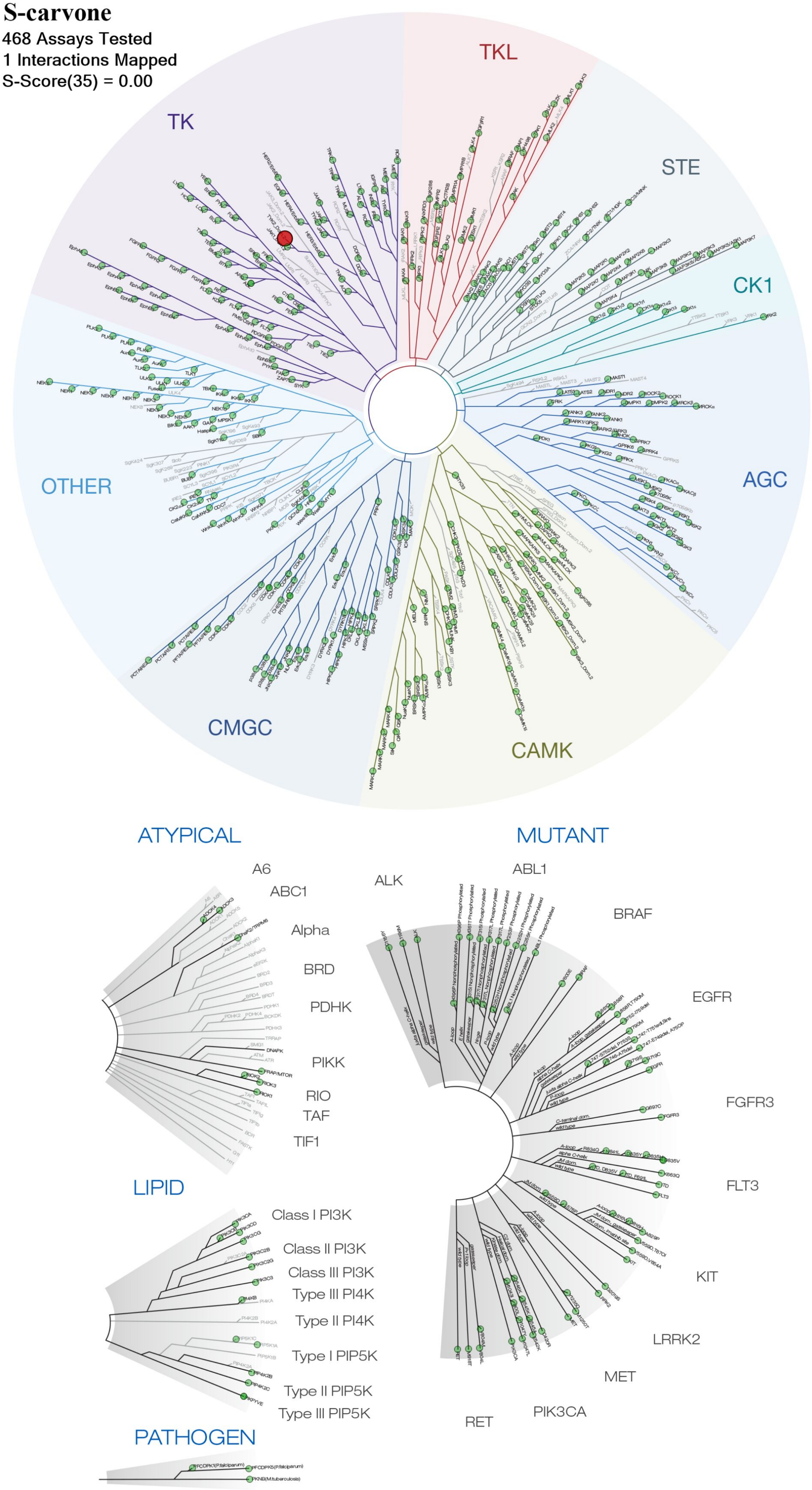

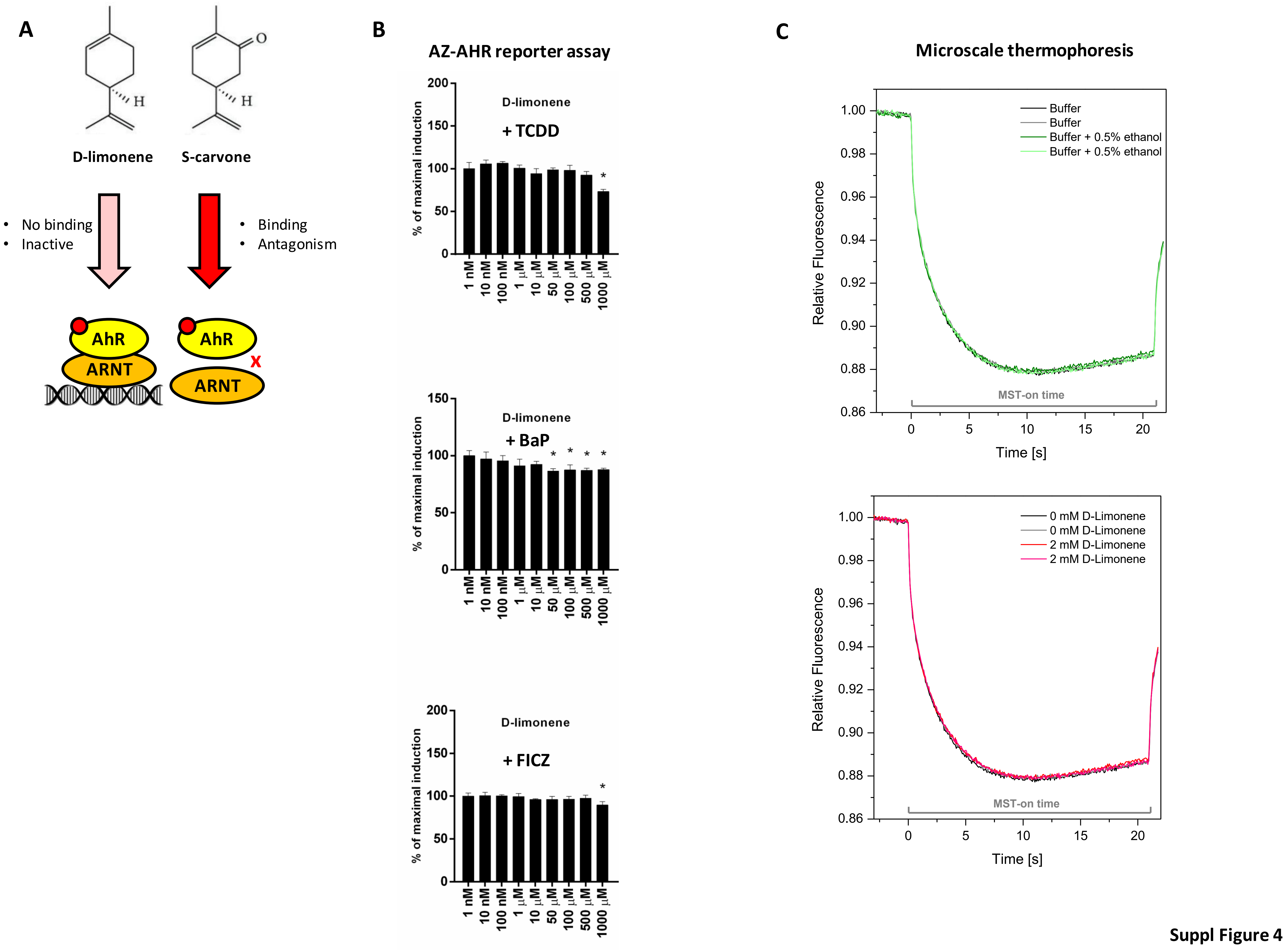


Suppl Figure 4

| % of inhibition | | R-CARVONE | | | | | S-CARVONE | | | | |
|---|---|---|---|---|---|---|---|---|---|---|---|
| | | 0 μM | 10 μM | 100 μM | 500 μM | 1000 μM | 0 μM | 10 μM | 100 μM | 500 μM | 1000 μM |
| **FICZ** | **100 nM** | 0% | 19% | 43% | 71% | 82% | 0% | 15% | 29% | 44% | 77% |
| | **1 μM** | 0% | 16% | 33% | 70% | 82% | 0% | 13% | 25% | 55% | 82% |
| | **10 μM** | 0% | 20% | 31% | 70% | 82% | 0% | 17% | 28% | 57% | 88% |
| | **50 μM** | 0% | 24% | 32% | 71% | 82% | 0% | 21% | 30% | 55% | 90% |
| | **100 μM** | 0% | 24% | 32% | 71% | 81% | 0% | 22% | 32% | 53% | 91% |
| | **200 μM** | 0% | 24% | 32% | 70% | 81% | 0% | 23% | 33% | 50% | 91% |
| **BaP** | **5 μM** | 0% | 24% | 63% | 94% | 97% | 0% | 18% | 45% | 89% | 95% |
| | **10 μM** | 0% | 16% | 61% | 95% | 97% | 0% | 19% | 44% | 92% | 97% |
| | **50 μM** | 0% | 19% | 49% | 93% | 97% | 0% | 19% | 37% | 90% | 96% |
| | **100 μM** | 0% | 20% | 45% | 91% | 96% | 0% | 19% | 33% | 87% | 95% |
| | **200 μM** | 0% | 21% | 43% | 87% | 94% | 0% | 18% | 31% | 84% | 93% |
| **TCDD** | **5 nM** | 0% | 12% | 56% | 98% | 100% | 0% | 39% | 60% | 93% | 100% |
| | **10 nM** | 0% | 23% | 59% | 96% | 100% | 0% | 14% | 55% | 89% | 99% |
| | **50 nM** | 0% | 2% | 29% | 80% | 97% | 0% | 8% | 24% | 62% | 97% |
| | **100 nM** | 0% | 3% | 26% | 69% | 88% | 0% | 8% | 20% | 53% | 90% |
| | **500 nM** | 0% | 4% | 25% | 62% | 87% | 0% | 8% | 19% | 46% | 88% |

**Suppl. Table 1. Reporter gene assay in AZ-AHR.** Inhibition of ligand-inducible AhR activity by carvones was calculated from plots shown in Figure 1C, as follows: % of inhibition at $c_X$ = 100*(1- (% of max. induction at $c_X$)/(% of max. induction at $c_0$))

### Nuclear translocation of AhR

| | | # of cells | # of fields of vision | # of AhR positive nuclei | % of AhR positive nuclei | Relative ligand efficiency (%) |
|---|---|---|---|---|---|---|
| **DMSO** | Exp #1 | 359 | 3 | 16 | 4.5 | n.a. |
| | Exp #2 | 438 | 4 | 6 | 1.3 | n.a. |
| **TCDD 20 nM** | Exp #1 | 550 | 5 | 235 | 43.1 | 100 |
| | Exp #2 | 453 | 4 | 198 | 43.6 | 100 |
| **BaP 7 µM** | Exp #1 | 446 | 4 | 143 | 31.9 | 100 |
| | Exp #2 | 486 | 4 | 158 | 32.6 | 100 |
| **FICZ 8 nM** | Exp #1 | 494 | 4 | 176 | 35.3 | 100 |
| | Exp #2 | 470 | 4 | 188 | 40.9 | 100 |
| **R-carvone 1 mM** | Exp #1 | 445 | 4 | 18 | 4.0 | n.a. |
| | Exp #2 | 429 | 4 | 2 | 0.5 | n.a. |
| **R-carvone + TCDD** | Exp #1 | 437 | 4 | 212 | 48.6 | 90 |
| | Exp #2 | 491 | 4 | 193 | 39.5 | 97 |
| **R-carvone + BaP** | Exp #1 | 454 | 4 | 144 | 32.0 | 101 |
| | Exp #2 | 396 | 4 | 169 | 42.3 | 107 |
| **R-carvone + FICZ** | Exp #1 | 481 | 4 | 191 | 40.0 | 109 |
| | Exp #2 | 496 | 5 | 191 | 38.7 | 102 |
| **S-carvone 1 mM** | Exp #1 | 570 | 5 | 23 | 4.0 | n.a. |
| | Exp #2 | 428 | 4 | 6 | 1.5 | n.a. |
| **S-carvone + TCDD** | Exp #1 | 485 | 4 | 233 | 48.6 | 99 |
| | Exp #2 | 462 | 4 | 157 | 34.3 | 79 |
| **S-carvone + BaP** | Exp #1 | 450 | 4 | 163 | 36.3 | 116 |
| | Exp #2 | 519 | 4 | 146 | 28.4 | 92 |
| **S-carvone + FICZ** | Exp #1 | 460 | 4 | 194 | 42.4 | 111 |
| | Exp #2 | 428 | 4 | 154 | 36.3 | 81 |

**Suppl. Table 2. Nuclear translocation of AhR – quantification of immunofluorescence.**
LS180 cells were treated for 90 min with S/R-carvones in combination with vehicle or AhR agonists TCDD, BaP and FICZ. Intracellular AhR was visualized with Alexa Fluor 488 labelled primary antibody. The whole staining protocol was performed in two independent experiments in technical duplicates. The AhR translocation was evaluated visually depending on the distinct signal intensity of AhR antibody in the nucleus and cytosol. For percentage calculation, approximately one hundred cells from at least four randomly selected fields of view in each replicate were used. Relative ligand efficiency was calculated as follows:

$$relative\ ligand\ efficiency = 100 \times \frac{(\#AhR\ positive\ nuclei)_{LIGAND+CARVONE} - (\#AhR\ positive\ nuclei)_{DMSO}}{(\#AhR\ positive\ nuclei)_{LIGAND} - (\#AhR\ positive\ nuclei)_{DMSO}}$$
